# Reconstructing the diet, trophic level, and migration pattern of Mysticete whales based on baleen isotopic composition

**DOI:** 10.1101/2020.10.04.301341

**Authors:** Philip M. Riekenberg, Jaime Camalich, Elisabeth Svensson, Lonneke L. IJsseldijk, Sophie M.J.M. Brasseur, Rob Witbaard, Mardik F. Leopold, Elisa Bravo Rebolledo, Jack J. Middelburg, Marcel T.J. van der Meer, Jaap S. Sinninghe Damsté, Stefan Schouten

## Abstract

Baleen from mysticete whales is a well-preserved proteinaceous material that can be used to identify migrations and feeding habits for species whose migration pathways are unknown. Analysis of δ^13^C and δ^15^N from bulk baleen has been used to infer migration patterns for individuals. However, this approach has fallen short of identifying migrations between regions as it is difficult to determine variations in isotopic shifts without temporal sampling of prey items. Here we apply analysis of δ^15^N values of amino acids to five baleen plates belonging to three species, revealing novel insights on trophic position, metabolic state, and migration between regions. Humpback and minke whales had higher reconstructed trophic levels than fin whales (3.4-3.5 versus 2.7-2.9, respectively) as expected due to different feeding specialization. Isotopic niche areas between baleen minima and maxima were well separated, indicating regional resource use for individuals during migration that aligned with isotopic gradients in Atlantic Ocean particulate organic matter. δ^15^N values from phenylalanine confirmed regional separation between the niche areas for two fin whales as migrations occurred and elevated glycine and threonine δ^15^N values revealed physiological changes due to fasting. Simultaneous resolution of trophic level and physiological changes allow for identification of regional migrations in mysticetes.

## Introduction

Mysticete whales are a concern for ecosystem based management after their populations were decimated by whaling (1, 2). Despite their current protected status, many details of the migratory patterns and feeding ecology of individual mysticetes remains uncertain and limited to a broader understanding of population scale patterns. This is especially true at the level of metapopulations and any additional information on ecological niche separation may help to inform policy makers to protect key habitats (3). New tools are now used to unravel migrations of mysticete whales, such as retrospective analysis of historical landing data (1, 4) and satellite tags to track both migratory paths and feeding strategies (5-9). However, tags provide only a small observational time window and are difficult to successfully deploy due to the mobility and lifestyle of these animals (10).

Feeding strategy and trophic ecology of mysticetes have primarily been identified through visual confirmation during feeding or stomach content analysis from strandings and historic catch data (11, 12). However, prey in the stomach of deceased animals only represent the animal’s last meal with the presence and the content (13) depending on the health status of the animal prior to death (i.e. trophic downgrading due to sickness). In contrast, proteinaceous materials that are continually produced across an animal’s lifetime (e.g. baleen, earplugs) provide a continuous record of metabolic processes (14, 15) and dietary composition across the time period that they have been produced and are thus useful in identifying prey composition and feeding strategies over a long period prior to death (16, 17). As a metabolically inert tissue, the incremental deposition of baleen faithfully records the dietary composition of the animal from when it is deposited until it is either lost or worn away. Baleen is composed almost entirely of keratin derived from metabolites of the bloodstream (18) and captures a long term record of the animal’s blood protein during keratin synthesis (19). This is in contrast to erythrocytes or skin tissue that provide a single integrated snapshot of isotopic composition across approximately one to two weeks to four months, respectively, depending on turnover within the pool of carbon (C) or nitrogen (N) being examined and the size of the animal (20). Whole lengths of baleen often reflect the dietary conditions from several months to several years depending on their growth rate and sampling distance from the gums.

The isotopic composition of C and N in baleen protein (expressed as δ^13^C and δ^15^N values, respectively) provides insights about the diet composition (10, 21-23). Typically, consumers have higher δ^13^C and δ^15^N values by 0.5-2‰ and 0.5-5‰, respectively compared to their diet in the system being examined (24, 25). These enrichments, or trophic discrimination factors (TDFs), vary with species, tissue type, metabolism, and diet quality and therefore require consideration of the available ecological context when assigning a TDF to a species within an ecosystem (26, 27). Dietary estimates reconstructed using bulk isotopic values reflect a mixture of influencing factors and often provide results that are inconclusive or muddled (28). This is especially true in systems where the isotopic baseline supporting production changes or the trophic level that animals feed at have shifted (29, 30). The δ^15^N values of resources supporting primary production can shift substantially, both spatially and temporally, depending on the balance of biogeochemical processes affecting available N sources (NO_3_^-^, NO_2_^-^ NH_4_^+^, N_2_) within an ecosystem (31-34). This occurs, for example, with wide ranging mysticetes in the north Atlantic Ocean where considerably higher δ^15^N values are observed for particulate organic matter (POM) in the higher latitudes (>70°N, 6 to 10‰) compared to lower mid-Atlantic areas (10°N, −1 to 1‰; (35). These shifts in baseline δ^15^N values are expected to interfere with estimates of trophic level for North Atlantic mysticetes as they migrate between mid-Atlantic breeding grounds and high-latitude feeding grounds. Correction for this baseline shift would usually require extensive sampling of primary consumers across both regions (24, 36).

The issue depicted above may be resolved by compound specific analysis of the δ^15^N values of amino acids contained in baleen. Isotopic differences that arise due to metabolic pathway differences between amino acid types can help to address regional shifts caused by the underlying δ^15^N baseline by providing simultaneous temporal information on trophic level and baseline δ^15^N values supporting the consumer (37-39). Baseline isotopic values are provided by source amino acids that undergo little change as they are metabolized (e.g. phenylalanine (Phe), methionine, tyrosine, and lysine; (40, 41). Additionally, trophic amino acids undergo considerable fractionation as they are metabolized (glutamic acid (Glu), aspartic acid, alanine, isoleucine, leucine, proline, and valine) and so-called ‘metabolic’ amino acids that undergo variable fractionations depending on the animal’s physiology or dietary composition (glycine (Gly), threonine (Thr)). Through the utilization of both source and trophic amino acids, a trophic discrimination factor (TDF) (38), and β, the difference between trophic and source amino acids in the underlying primary producers, trophic level estimates for individuals can be calculated. Trophic level estimates inherently integrate underlying baseline shifts that have occurred during migrations between breeding and feeding grounds. Uncertainties remain about the effects of diet quality and metabolic effects associated with routing of compounds (i.e. higher fractionation associated with poorer assimilation efficiency) or excretion pathways (i.e. excretion of urea versus ammonia), but these are incorporated into the high level of uncertainty assigned to the TDF (7.6±1.5‰; (40) and resulting trophic level estimates.

In this study we are among the first to combine bulk stable isotope with analysis of δ^15^N in amino acids from baleen sourced from stranded or bowcaught fin whales (n=3, *Balaenoptera physalus*), a stranded humpback whale (*Megaptera novaeangliae*), and a minke whale (*Balaenoptera acutorostrata*), all opportunistically sampled in the Netherlands. δ^15^N values of amino acids from baleen allows for resolution of trophic effects from changing regional δ^15^N values and characterization of potential metabolic effects from fasting and episodic feeding. Additionally, we provide evidence of feeding in different regions due to migration between Mid-Atlantic breeding and high-latitude feeding areas through application of isotopic niche areas from trophic level corrected bulk isotope data from baleen.

## Methods

### Sample collection

Baleen plates were collected from dead mysticetes that stranded on the Dutch coast or brought into Dutch harbors caught on a ship’s bow in 2012 and 2013 (Table 1). Baleen from three fin whales were acquired from the faculty of Veterinary Medicine of Utrecht University, the baleen from the humpback whale (*Megaptera novaeangliae*) was acquired from Naturalis Biodiversity Center, Leiden, and the minke whale (*Balaenoptera acutorostrata*) baleen was acquired from a private collection.

**Table 1:**
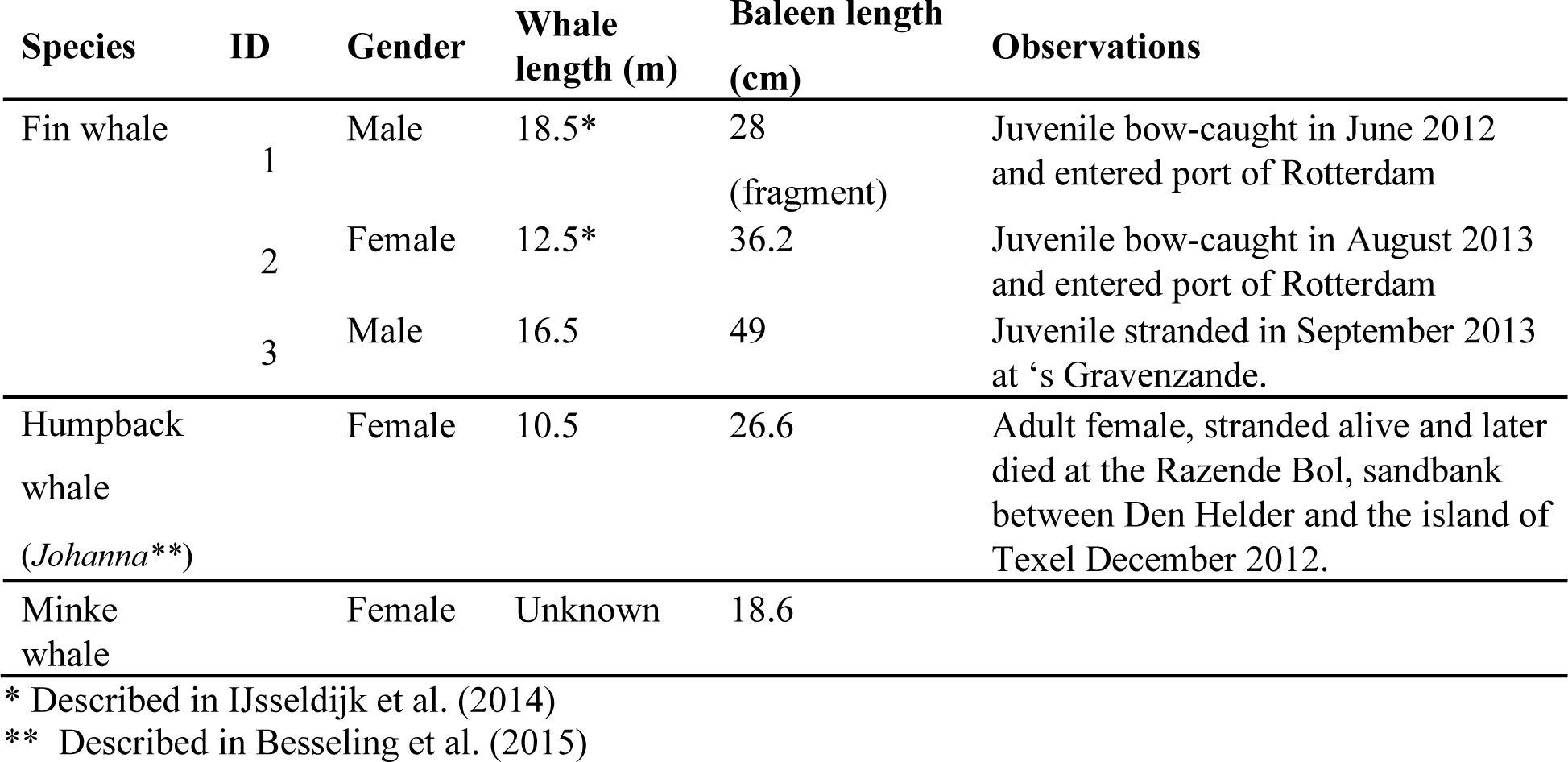
Information on study specimens and baleen.

### Sample preparation

One baleen plate per individual whale was air dried (>2 days), cleaned with bidistilled water, then dichloromethane, and dried at 40°C for ∼10 hours. Powdered keratin was collected using a hand drill (3 mm bit) along the leg (labial side) of the plate at either 0.5 cm or 1 cm intervals (representing a range of ∼2 to 4 weeks of growth between samplings) depending on the relative size of the plate, from the gingiva along the full length of the main plate. Powdered baleen was collected and stored at −20°C until further processing. Since baleen grows continuously throughout a whale’s life the material closest to the gingiva reflects the most recently produced layer with the material farther away from the gums reflecting increasingly older periods of the whale’s foraging history. Subsampled material (3-5 mg) for analysis of δ^15^N of amino acids were selected based on the variation within the bulk δ^15^N values observed for each individual and target minimum and maximum values observed across the lengths of plate.

### Bulk stable isotope analysis

Approximately 0.5–0.8 mg of dry, homogenized keratin powder was weighed into tin cups in duplicates for determination of carbon (δ^13^C) and nitrogen (δ^15^N) isotopic ratios for bulk material, as well as carbon and nitrogen content (%) of bulk biomass. Samples were analyzed on a Flash 200 elemental analyzer coupled to a Delta V Advantage isotope ratio mass spectrometry (Thermo scientific, Bremen).

Stable isotope ratios are expressed using the δ notation in units per mil:

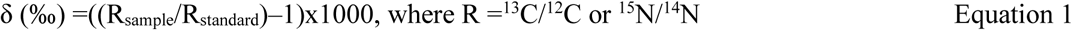

and expressed versus Vienna Pee Dee Belemnite (VPDB) for δ^13^C and atmospheric N_2_ (air) for δ^15^N. A laboratory acetanilide standard with δ^13^C and δ^15^N values calibrated against NBS-22 and IAEA-N1, respectively, and known %TOC and %TN contents, was used for calibration. Analytical precision for the standards (urea, casein) for δ^13^C and δ^15^N analyses were 0.18‰ and 0.2‰, respectively.

### Amino acid sample preparation

The method is a modified version of the amino acid analysis method by Chikaraishi et al. (2007) as described in Riekenberg, van der Meer (42). In short, samples were hydrolyzed, derivatized into N-pivaloyl/isopropyl (NPiP) derivatives and analyzed in duplicate with a Trace 1310 gas chromatograph coupled to a Delta V Advantage isotope ratio mass spectrometer (Thermo Scientific, Bremen) via an IsoLink II. Details about the temperature ramp, program settings, and normalization procedures are provided in (42). We report δ^15^N values for 12 amino acids including alanine, aspartic acid, Glu, Gly, isoleucine, leucine, lysine, Phe, serine, Thr, tyrosine, and valine. The precision for samples and standards was <0.5‰ for all amino acids across the 14 sequences that comprise this data set.

### Estimating growth intervals

To examine the relative rates of change for the oscillations in the δ^15^N values for bulk material along the length of the main plate, we fitted a generalized additive model (GAM) for each individual. GAM models were produced using the geom_smooth function in the ggplot2 package with model= “gam” to apply smoothing parameters selected by data driven methods using Akaike information criteria to time series in R (v4.0.0) with R Studio (v1.1.463) (17, 43). Oscillations within δ^13^C values for individuals were less distinct, having a smaller range than those for δ^15^N values often due to closer similarity in prey δ^13^C values and are known to be further confounded due to coastal foraging in areas with gradients in δ^13^C (17). Therefore, δ^15^N values were used to estimate baleen growth rates for each individual by assuming the oscillation between sequential δ^15^N value minima along the baleen record represented migratory annual movements between foraging grounds. Growth estimates were determined as the distance between sequential minimum δ^15^N values and this interval was used to estimate a weekly growth rate as in Busquets-Vass, Newsome (17). To further clarify the midpoint between minimum and maximum periods for δ^15^N values we plotted a linear regression across δ^15^N values for each baleen and binned regions of each baleen into minimum (below midpoint) and maximum (above midpoint) values depending on relative position to the conditional mean to allow for further analysis of regional differences for δ^15^N values.

### Trophic level calculations

Trophic level estimated from baleen amino acids are presented using weighted averages for both source and trophic amino acids presented in Chikaraishi, Ogawa (40) modified with a tissue specific TDF previously estimated for blue whales (*Balaenoptera musculus*) in Busquets-Vass, Newsome (17).

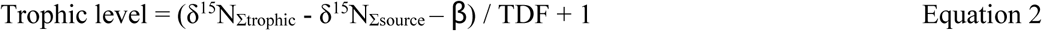

where δ^15^N_Σtrophic_ and δ^15^N_Σsource_ are the weighted mean values for grouped trophic (alanine, aspartic acid, glutamic acid, isoleucine, leucine, and valine) and source (lysine and phenylalanine) amino acids, respectively, β, the ‰ difference between Glu and Phe in the underlying phytoplankton is assumed to be 3.4‰, and a TDF of 4‰ accounting for the discrimination in trophic amino acids between the baleen whales and their diet. The TDF of 4‰ has been scaled from the 1.8‰ trophic enrichment factor estimated for δ^15^N for bulk material from *B. musculus* baleen (17) assuming scaling factors of 3.4‰ and 7.6‰ for trophic discrimination factors for bulk and amino acid trophic level determination, respectively. Error propagation for trophic level gives a standard deviation of ±0.8 using the equation presented in Ohkouchi, Chikaraishi (39).

Correction for trophic enrichment to establish baseline estimates for δ^15^N using phenylalanine were calculated as

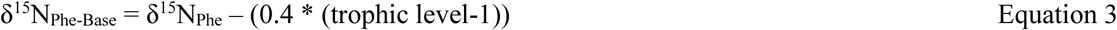

where 0.4‰ is the small enrichment observed for Phe during metabolism (40) and trophic level calculated for each individual (Table 1).

Trophic level estimates were further used to estimate baseline δ^13^C and δ^15^N values for bulk measurements using the equations:

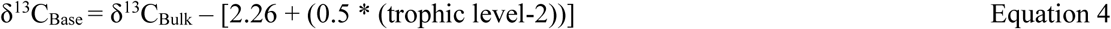

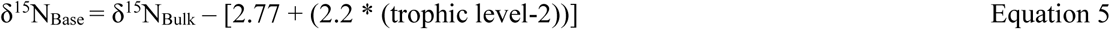

where δ^13^C_Bulk_ and δ^15^N_Bulk_ represent the C and N isotopic composition of bulk material, 2.26‰ and 2.77‰ represent the offset between diet and baleen for carbon and nitrogen (44), 0.5‰ and 2.2‰ represent the offsets for trophic enrichment for carbon and nitrogen for the trophic levels supporting the whales prey (25), and trophic level is the average estimate for each individual (Table 1). Wilcoxon signed rank t-tests were used to examine individual amino acid δ^15^N values between regions of baleen.

### Niche modeling

To analyze differences in isotopic niches within each individual baleen, standard ellipse areas corrected for their sample size (SEA_c_) were constructed containing 70% of the variation in each group for the binned minimum and maximum values for δ^13^C_Base_ vs δ^15^N_Base_ for each individual using the SIBER package (45). The overlap between groups was characterized through calculation of the Euclidean distance between the centroids for both minimum and maximum SEA_c_, followed by a residual permutation and Hotelling T^2^-test to evaluate statistical differences (46, 47) between the areal coverage of the two niches (α=0.05) using the package ‘Hotelling’.

## Results

### Bulk δ^13^C and δ^15^N values

The δ^13^C values for all individuals fell within the range of −17.5 to −20 ‰ across all baleens, with oscillations of 0.5 to 1.5‰ that generally mirrored changes observed in δ^15^N values, with some deviations (Fig. 1, Supplemental Table 1). δ^13^C values for the fin whales were similar among individuals and higher (−18.9 to −19.2‰) than for the humpback whale (−19.6‰), but lower than for the minke whale (−18.1‰; one-way ANOVA: F_4,252_=77, *p*<0.001). Within-individual variation in δ^15^N values was larger than seen for δ^13^C values (maximum within-individual range in δ^15^N is 11.2 to 14.8‰ in the humpback whale). Oscillations in δ^15^N values also showed greater amplitude from 0.5 to ∼3‰, with median values for the humpback and minke plates (12.8‰ and 12.2‰) higher than those for all three fin whale plates (9.3 to 10‰: Supplementary Table 1). δ^15^N values were higher for both the minke and humpback whale (11.8 and 12.8‰, respectively; one-way ANOVA: F_4,252_=238, *p*<0.001) than for the fin whales (9.2 to 10‰).

**Figure 1:**
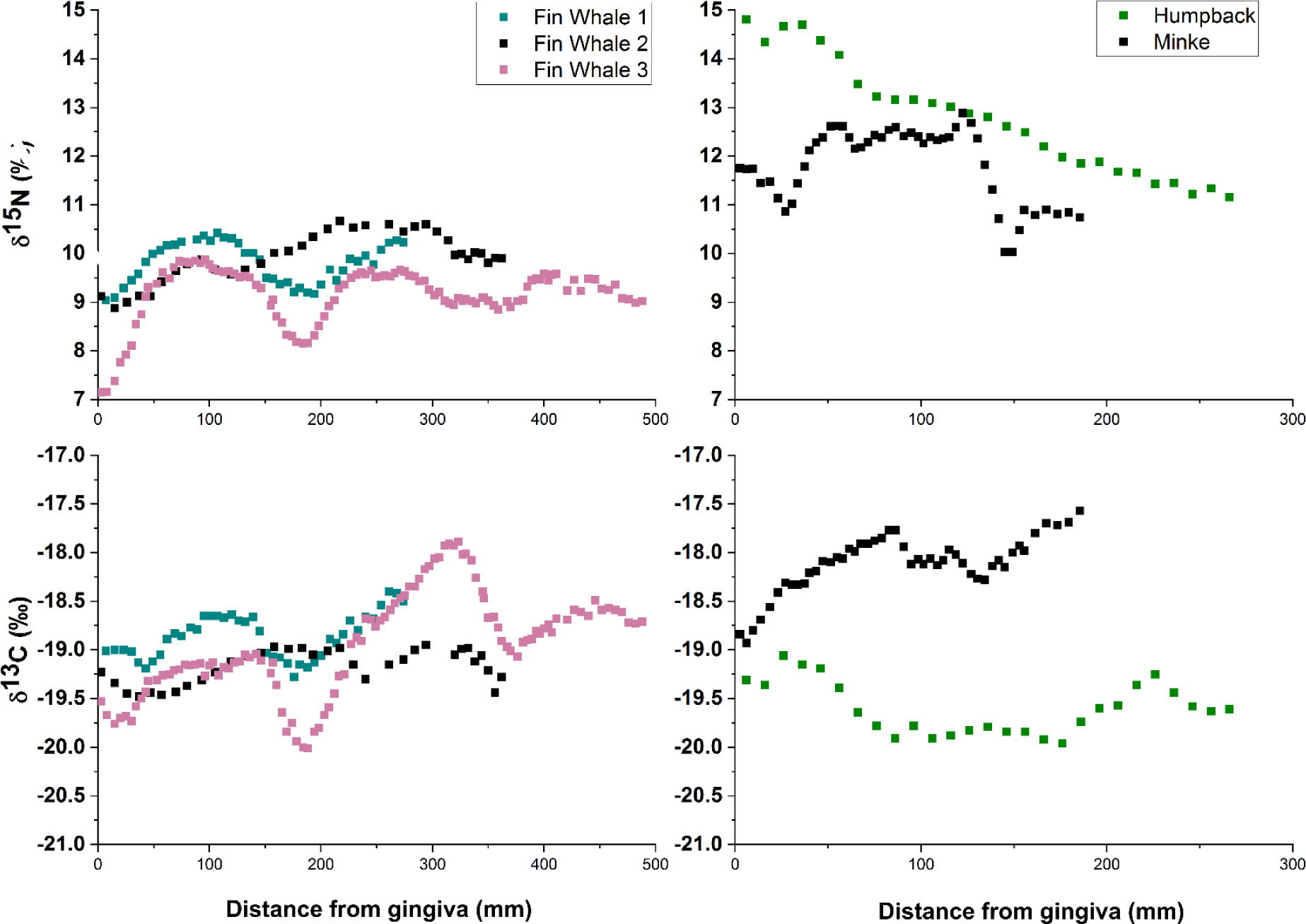
Bulk δ^15^N and δ^13^C values from incrementally sampled baleen plates of 5 individual mysticetes originating from the North Atlantic.

The three fin whales (FW1, FW2, FW3, Fig. 2) displayed regular oscillations in δ^15^N values that imply baleen growth rates of 2-3.5 mm week^-1^, calculated based on GAM modelling (Table 2), following the approach of Aguilar, Giménez (48). The minke whale (MN, Fig. 2) showed less regular minima that corresponded to a growth rate of 2.3 mm week^-1^, while the humpback whale displayed no distinct δ^15^N minima but rather a continuous decrease in δ^15^N value from the gingiva across the full length of baleen (from 14.8 to 11.6‰; HB, Fig. 2) with slight oscillations from which no growth estimate could be reasonably estimated. Linear regressions applied to the δ^15^N values for bulk baleen indicated regions in the baleen that were above (black) and below (gray) the conditional mean (Fig. 2). The minimum and maximum periods of these oscillations reflect the net effects from metabolism, trophic position, and the underlying values of the resources being utilized during each individual’s migrations. Minimum and maximum periods for δ^15^N values (grey and black bars, Fig. 2) are thought to reflect residence times in different waters within the Atlantic and Arctic Oceans, with the differences in amplitudes of oscillations reflecting the net effects from different migrations that occurred within an individual’s lifetime and the seasonal decrease in the excretion of ^15^N in urine as fasting occurs (48, 49). These minimum and maximum periods were utilized to target amino acid δ^15^N samples (hash marks Fig. 2) in order to maximize the potential differences between samplings in each period.

**Table 2:** Trophic level (TL) means for each individual. TL is a unitless measurement, n represents the number of amino acid measurements along the length of baleen for each individual.

| Individual | SD | TL | n |
| --- | --- | --- | --- |
| Fin Whale 1 | 0.4 | 2.7 | 10 |
| Fin Whale 2 | 0.3 | 2.7 | 11 |
| Fin Whale 3 | 0.5 | 2.9 | 16 |
| Humpback | 0.3 | 3.4 | 12 |
| Minke | 0.4 | 3.5 | 11 |

**Table 3:** Baleen growth estimates. GAM results assessing the fluctuations of δ^15^N in baleen plates and the resulting growth estimates from these models.

| Individual | n | E.D.f. | F | Adjusted $R^2$ | P | Deviance explained (%) | $\delta^{15}\text{N}$ minima interval (cm) | Weekly growth interval (mm) |
| --- | --- | --- | --- | --- | --- | --- | --- | --- |
| Fin Whale 1 | 42 | 8.6 | 82 | 0.95 | < 0.001 | 95.6 | 15.6 | 3 |
| Fin Whale 2 | 35 | 8.7 | 92 | 0.96 | < 0.001 | 96.9 | 10.4 | 2 |
| Fin Whale 3 | 105 | 8.8 | 112 | 0.91 | < 0.001 | 91.4 | 13.5, 16.3, 18.1 | 2.6-3.5 |
| Humpback | 27 | 3 | 279 | 0.98 | < 0.001 | 97.8 | NA | NA |
| Minke | 48 | 8.4 | 39 | 0.88 | < 0.001 | 90.2 | 12.2 | 2.3 |

**Figure 2:**
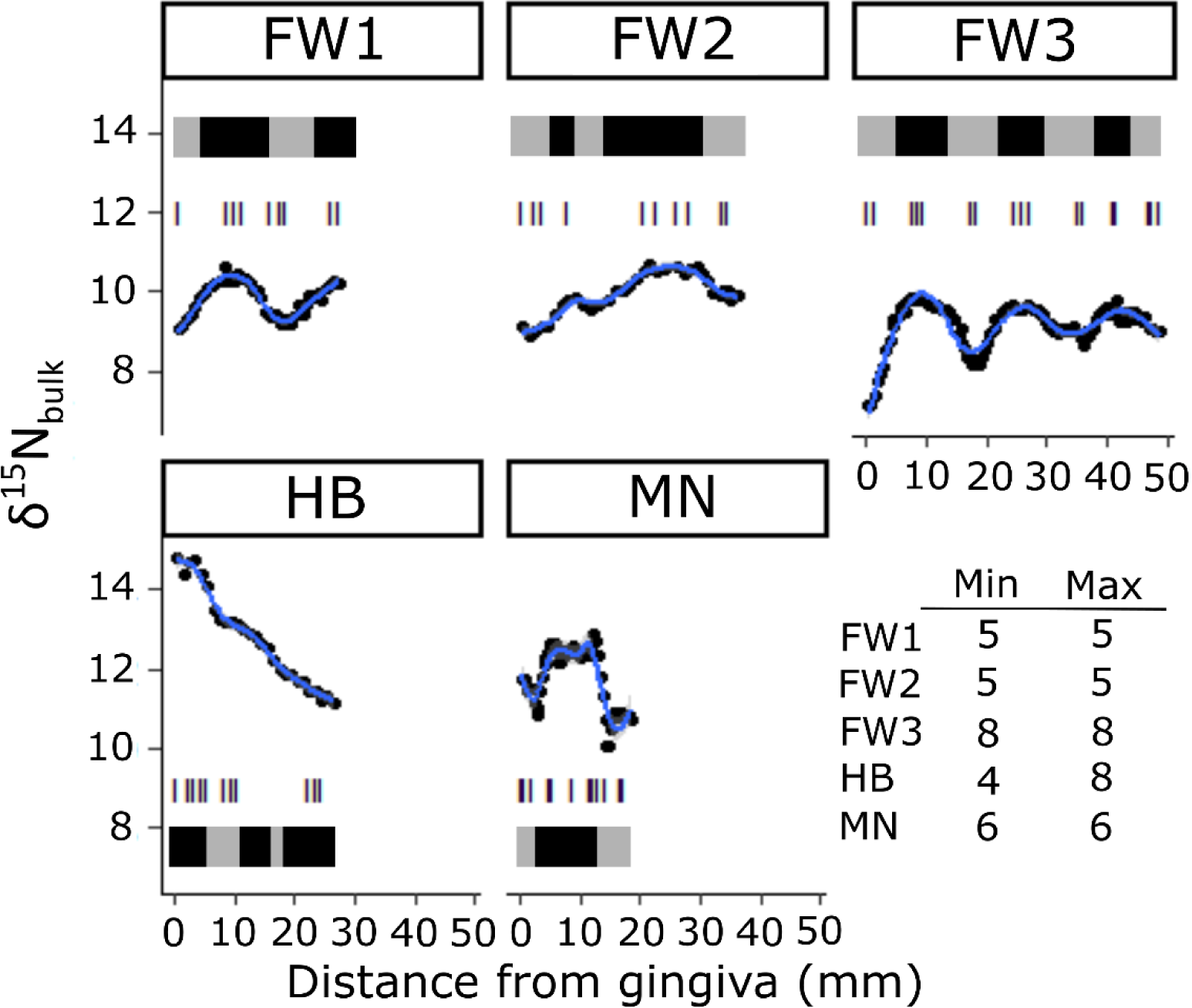
GAM models fit to baleen δ^15^N_bulk_ values to identify minimum and maximum periods for δ^15^N values in baleen for each individual (blue lines). Black shaded regions indicate periods when δ^15^N values were above (Max) and grey shaded regions indicate periods when δ^15^N values were below (Min) the conditional mean for a linear regression applied to bulk δ^15^N values. Hash marks indicate sampling intervals for determining δ^15^N values for amino acids and the table in the bottom right panel indicates the samples located in Min and Max periods for each individual whale.

### δ^15^N of amino acids

Trophic levels were estimated using the weighted means of the δ^15^N values for trophic and source amino acids (see methods Eq. 2) with means ranging from 2.7 to 2.9 for the fin whales, 3.4 for the humpback whale, and 3.5 for the minke whale (Fig. 3A). The trophic level estimates did not vary significantly (two-way ANOVA, all *p* >0.05) between the minimum and maximum periods examined along the baleen. Therefore, single trophic level estimates from the average for each individual whale were utilized for trophic level corrections applied in the calculations for baseline corrected δ^13^C and δ^15^N values (δ^13^C_Base_ and δ^15^N_Base_; Eq 2; Table 2). This correction is also applied to Phe to reduce δ^15^N values representing the calculated value supporting the base of the food web (δ^15^N_Phe-Base_) by correcting for fractionation due to trophic levels using the measured TL estimate (Eq. 3). Changes in the δ^15^N_Phe-Base_ values reflect the differences in underlying regional N source values supporting each individual during the period when the plate was formed. The δ^15^N_Phe-Base_ values for time intervals with minimum and maximum bulk δ^15^N values were found to be significantly different between individuals and between minimum and maximum δ^15^N value periods (two-way ANOVA: Individuals, F_4,54_=5.3, *p*=0.001; min/max F_1,54_=11.4, *p*=0.001; Interaction: F_5,54_ =6.9, *p*<0.001; Fig 3b). FW3 was found to have higher δ^15^N_Phe-Base_ in the maximum periods for bulk δ^15^N values than in the minimum periods (Wilcoxon ranked t-test, Z_8_= −2.5, *p*= 0.008). While for FW2 the δ^15^N_Phe-Base_ values were also higher in the maximum bulk ^15^N period than in the minimum periods (Z_5_=-1.9, *p*= 0.06), but not statistically significant using a threshold of α=0.05 (Fig 3b).

**Figure 3:**
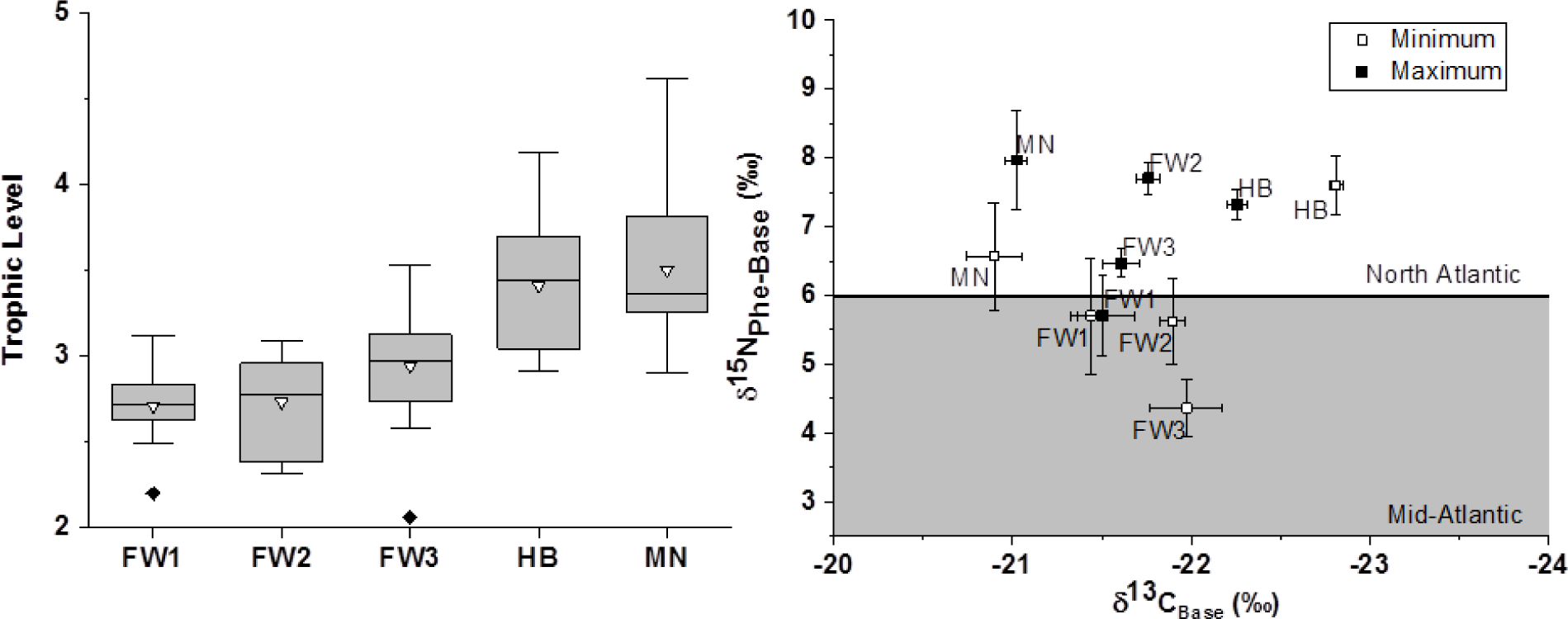
A) Trophic level estimates for each individual and B) Baleen baseline mean δ^13^C_Base_ and δ^15^N_Phe-Base_ values minimum and maximum values across the lengths of baleen for each individual; mean ± SE. The grey shaded area indicates the δ^15^N baseline isotope value for Mid-Atlantic (2 to 6.5‰) versus the North Atlantic (6 to 10‰) oceans (Magozzi et al 2017; Trueman et al. 2019).

Thr and Gly are both metabolic amino acids that provide indications for diet composition and fasting state of the individuals, respectively (50, 51). δ^15^N values of Phe were subtracted from both Thr and Gly δ^15^N values to correct metabolic AA δ^15^N values for changes resulting from baseline values (Fig 4a & b). δ^15^N_Thr_ values were not significantly different when examined between minimum and maximum periods for bulk δ^15^N values (two-way ANOVA, all *p*> 0.05), but were found to be generally higher for the fin whales (one-way ANOVA: F_4,53_ = 6.8, *p*<0.001; Fig. 4a) with a wider range (−29.3 to −11.2‰) than for the humpback whale (−28.4 to −25.9‰) and minke whale (−32.3 to −21.6‰). δ^15^N_Gly_ values were significantly different between minimum and maximum periods for individuals (two-way ANOVA: Individuals F4,50= 112, *p*<0.001; min/max F_1,50_= 23.9, *p*<0.001). For all individuals, the means for the minimum periods were higher than for the maximum, but was only statistically higher for FW3 (Z_8_=2, *p*=0.04), although FW1 (Z_8_=1.9, *p*=0.06) was close to being significant at a threshold of α=0.05 (Fig.4b).

**Figure 4:**
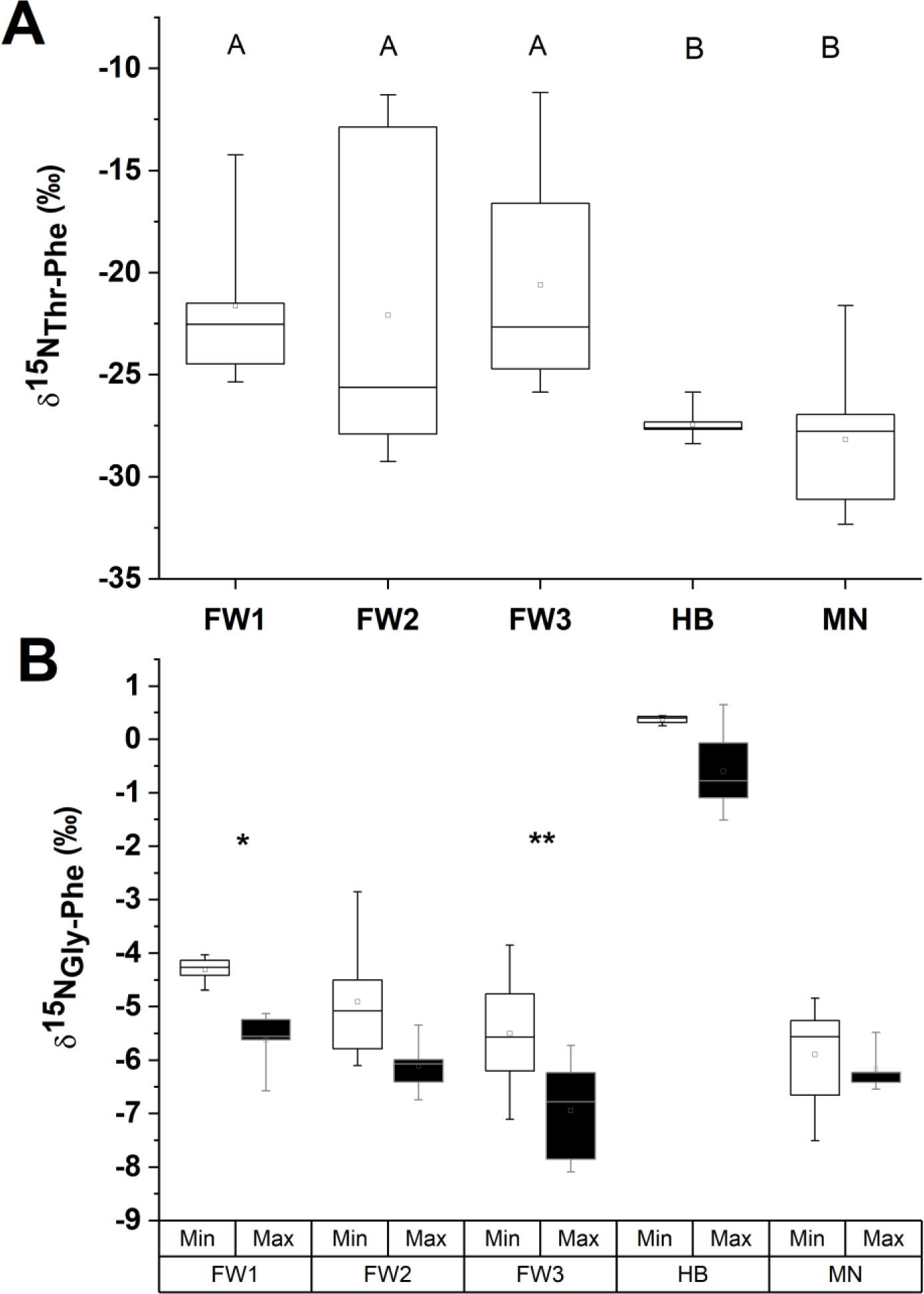
δ^15^N for A) threonine and B) glycine for the five mysticete individuals to assess possible starvation and fasting effects between individuals and baleen periods, respectively. Both Thr and Gly have had been corrected against Phe to remove underlying source AA variation. Letters indicate significant differences as indicated by a post-hoc Tukey’s test (α=0.05). For Gly, Wilcoxon ranked t-tests were used to compare between baleen regions for each individual. * indicates *p =* 0.06 and ** indicates *p* < 0.05.

### Isotopic niche overlap within individuals

Isotopic niche areas (SEAc) were calculated by applying a Bayesian statistical model (SIBER package) to the larger data set of trophic level corrected δ^13^C_Base_ and δ^15^N_Base_ values (Eqs. 4 & 5). The overlap between the minimum and maximum baleen periods ranged from 0 (implying different values for basal resources) to 56% (indicating some overlap) with Hotelling’s T^2^-test, indicating significant differences between periods for all individuals (Fig. 5). Separation of SEAc areas for periods predominately occurred along the y axis (δ^15^N_Base_) for all individuals besides HB, where separation occurred across the x axis (δ^13^C_Base_).

**Figure 5:**
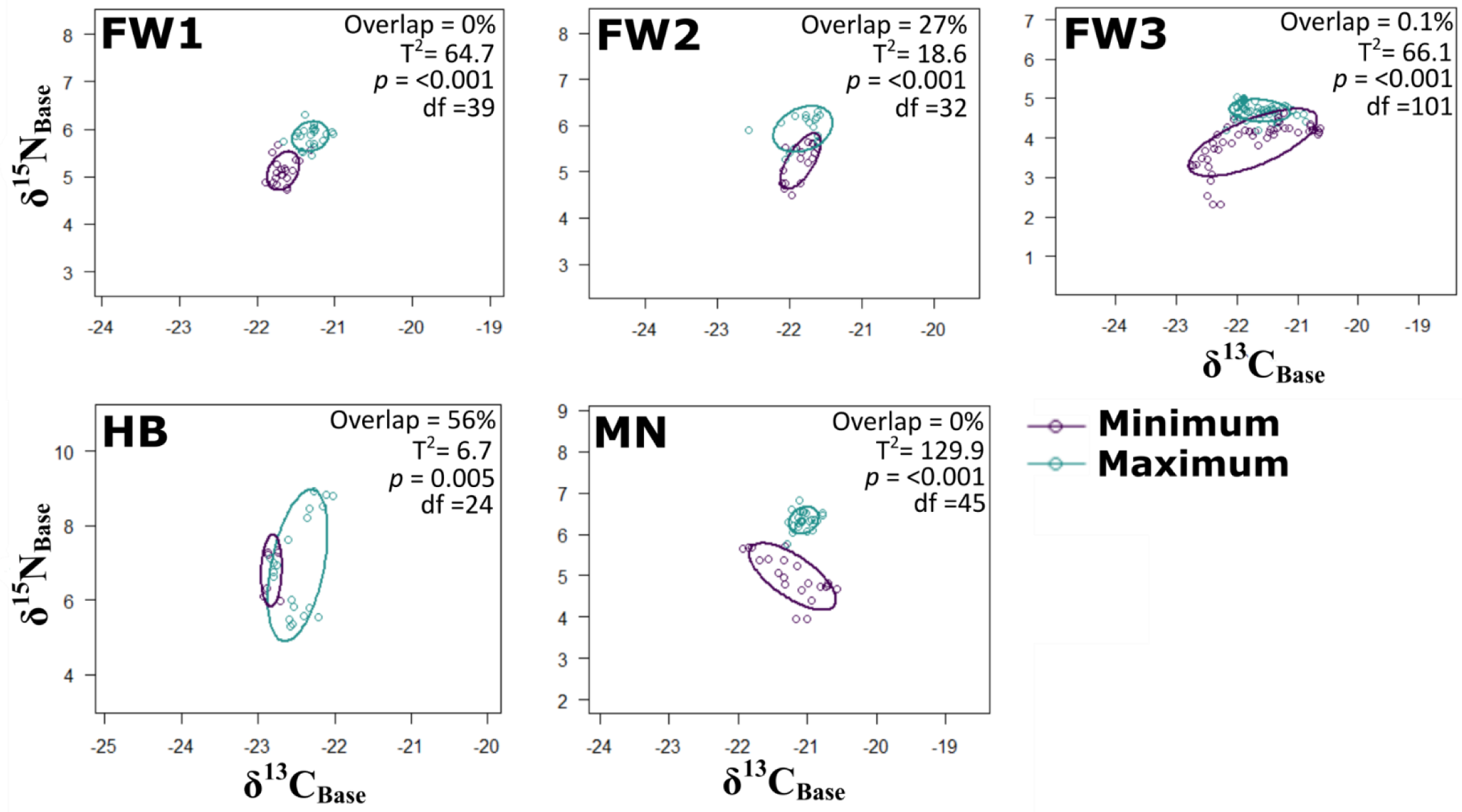
Standard ellipse areas corrected for sample size for baleen regions pooled into minimum and maximum periods as determined by GAM models on δ^15^N values for each individual. Overlap is the percentage of ‰^2^ areal overlap between the periods. Significant *p* values from Hotelling’s T^2^-test indicate where baleen values occupy different isotopic niches due to feeding in regions with distinct isotope resource values.

## Discussion

### Growth Rates for Baleen

The marked oscillations in δ^15^N values of baleen are assumed to reflect residence times in Mid-Atlantic breeding grounds (minima) and high-latitude feeding grounds (maxima) with substantial differences in the underlying δ^15^N values for POM in these regions (−1 to 1‰ versus 6-10‰, respectively) and differences in excretion of δ^15^N depending on fasting status during migrations (48, 49). The changes in amplitude of the oscillations in baleen reflect resource use and feeding status during migrations that occurred within an individual’s lifetime. The humpback whale did not display readily apparent oscillations in δ^15^N values, but rather a distinct increase in δ^15^N value that progressed along the length of baleen, suggesting different migration behavior than observed for the other whales in this study. The estimates for baleen growth rates for the 4 individuals with recurring minima in δ^15^N_bulk_ values (FW1-3 & MN; 10.4-16.3 cm y^-1^; Table 1) agree with previous estimates from blue (17), minke (12.9 cm y^-1^, (52)) and bowhead whales (*Balaena mysticetus*; 16-25 cm y^-1^, (19)), all of which assume continuous growth across seasons. The length of plates in this study are relatively short due to the availability of stranded and bow-caught animals examined and represent a maximum of 3 years of migration (FW3, with 4 δ^15^N minima) within these individuals. Other studies have examined considerably longer plates (17, 53, 54).

### Trophic Levels

Trophic level estimates from minimum and maximum regions of δ^15^N values along the baleen are expected to reflect the largest contrasts between food resources throughout the multi-year periods recorded in the baleen. Trophic level estimates from amino acid δ^15^N values indicated no significant changes in TLs across the baleen records for these 5 individuals. Consistent trophic levels throughout migratory periods reflect a more-or-less continuous utilization of the same kind of prey, without considerable periods of specialization, e.g. a switching between fish and krill during migration. The reconstructed trophic levels were significantly lower (one-way ANOVA: F_4,55_=11.7, *p*<0.001) for the fin whales than for the humpback and minke whales. This is fully consistent with the fact that fin whales preferentially consume krill in areas where these are abundant and only occasionally feeding opportunistically on schools of small fish when krill is scarce (55, 56). The higher reconstructed trophic levels for both humpback and minke whales are expected as their foraging typically includes larger number of small fishes less than 30 cm length such as herring (*Clupea harengus*) and sprat (*Sprattus sprattus*) (57-59). Trophic level estimates observed in the minimum and maximum regions of δ^15^N values along the baleen are expected to reflect the largest contrasts between food resources throughout the multi-year periods recorded in the plate. Estimates calculated from amino acid δ^15^N values indicated no significant changes in trophic levels across the baleen records for these 5 individuals. Consistent trophic levels throughout migratory periods reflects a more or less continuous utilization of the same prey, without considerable periods of specialization or switching between smaller fishes and krill during migration and no major effect of seasonal variations in ^15^N excretion rates. Trophic level estimates for all of the whales (ranging from 2.7 to 3.5) were based on a relatively small TDF (4‰) compared to the typically applied 7.6‰ value (the difference between the amino acids Glu and Phe in marine organisms, ((40)) for TDF (38, 41), but yield comparable trophic levels as determined through stomach contents for other mysticete whales (60). The small TDF reflects the increased similarity between the protein quality of the diet and the consumer, resulting in less reworking and therefore less fractionation of amino acid N during metabolism (38, 61).

### Metabolism of whales

The amino acids Gly and Thr have been found to respond to fasting and protein deficiency in mammals through variable enrichments in δ^15^N values for each AA (50, 51) as metabolism of stored lipid resources occurs. The δ^15^N values of glycogenic amino acids (Gly, serine, proline, and aspartic acid) are expected to increase as exogenous material becomes limited and endogenous protein starts to be metabolized (62). Fasting has been previously thought to occur as whales migrate southward out of their northern feeding grounds towards breeding sites (63) and feeding becomes more confined to opportunistic feeding events (48, 53). Baseline-corrected δ^15^N_Gly_ values were statistically higher in the minimum periods for bulk δ^15^N values than in the maximum periods for bulk δ^15^N in FW3 (Fig. 4b), although all five individuals had higher averages for minimum periods ranging from 0.3 to 1.4‰ and the difference for FW2 was nearly statistically significant (*p*=0.06). These isotopic enrichments for δ^15^N are lower than the observed shift (2-6‰) in glycogenic amino acids for fasting southern elephant seals (*Mirounga leonina*, Lübcker, Whiteman (50)). This suggests that the impacts from fasting on glycogenic δ^15^N values for mysticetes may be more limited than during breeding and molting events in other marine mammals. This smaller observed effect may be due to either the considerable body size and lipid stores or through subsistence with opportunistic feeding offsetting more extreme fasting effects (64). Although limited in scale, isotopic enrichment of the δ^15^N_Gly_ values onset in the same manner (during minimum periods) across all individuals regardless of species (Fig. 4b) and suggests that there is a fasting effect that occurs as fat stores are accessed during migrations towards breeding grounds when feeding becomes opportunistic.

Higher δ^15^N_Thr_ values in mammals have been observed to coincide with reduced protein quality in their diet causing reduced reverse fractionation with higher δ^15^N_Thr_ values indicating periods of potential starvation (51) although this mechanism is incompletely characterized. Extremely low values for δ^15^N_Thr_ are typical of marine mammals from higher trophic levels (65). Both humpback and minke whales have patterns of consistently low values (−27.4 ± 0.7‰ and −28.2 ± 3.2‰, respectively; Fig 4a) that are expected for adequate protein availability throughout the period of baleen examined. However, the fin whales (FW1-3) had higher δ^15^N_Thr_ values (−22.1 to −20.6‰) and larger ranges (3.7‰ to 7.4‰) due to higher δ^15^N values (−15 to −11‰) for Thr occurring intermittently throughout the baleen records. These higher values are potentially the result of protein deficiency and may mark protein deficiency or starvation events across an individual’s lifetime. This finding conflicts with decreased δ^15^N_Thr_ values observed in elephant seal whiskers during fasting (50), suggesting that there may be multiple effects impacting the metabolism of Thr during fasts that are more or less severe and warrant further investigation. There was no strong correlation between minimum and maximum periods for bulk δ^15^N values in the baleen and differences in δ^15^N_Thr_ suggest more episodic onset than the more regularly occurring fasting effects observed for δ^15^N_Gly_. Given these observations, the fin whales appear to have been more regularly under food stress than either the humpback or minke whale individuals examined in this study.

### Resource utilization

Minimum and maximum periods in baleen bulk δ^15^N values had distinct values for both δ^15^N_Phe-Base_ (Fig. 3B) and the isotope niches formed using the wider data set of δ^13^C_Base_ and δ^15^N_Base_ (eq. 4 & 5; Fig. 5). Distinct differences between the minimum and maximum bulk δ^15^N periods likely reflect underlying isotopic differences in resource values supporting the food web between Mid-Atlantic breeding grounds and high-latitude feeding grounds. Although few individuals have been tracked, all three species have been observed to make southerly winter migrations away from high-latitude feeding grounds (>70°N) with North Atlantic fin whales having been observed to migrate to the Azores (8), humpback whales as far south as the Antillean islands (66), and minke whales observed off the east coast of Florida (67). These large geographic separations between breeding and feeding grounds coincide with distinct underlying isotopic values for POM in those regions. Isoscapes, i.e. geographic maps of the underlying yearly averages for regional isotopic values of carbon and nitrogen (35, 68), characterize isotopic ranges for the POM sampled from both the mid-Atlantic (δ^13^C −28 to −30‰; δ^15^N −1 to 1‰) and high-latitude Arctic (δ^13^C −24 to −20‰; δ^15^N 6 to 10‰). The δ^13^C and δ^15^N values from these regions vary depending on local biogeochemical processes (e.g. lower δ^15^N values associated with oligotrophic conditions) and serve as variable end members for the source amino acids incorporated into baleen as it is produced from bloodstream metabolites derived from the animal’s diet. The variations in those values are likely to be dampened and reflect the relative feeding intensity (opportunistic feeding in transit), variations in seasonal values for underlying biogeochemical processes, and the relative turnover of the internal source AA pool during migration and breeding. The ranges in δ^15^N_Phe-Base_ (1.8 to 9.9‰), δ^13^C_Base_ (−20.6 to −22.9‰), and δ^15^N_Base_ (2.3 to 8.9‰) values all fall within the ranges expected for dietary intake of resources sourced from regions with distinct baseline isoscape values.

The larger ranges observed for δ^15^N_Phe_ and δ^15^N_Base_ (8.1 and 6.6, respectively) versus δ^13^C_Base_ (2.3‰) are likely due to increased turnover in N when compared to C within tissues as incidental feeding occurs (Trueman et al. 2019). N from incidental feeding is likely to be directly metabolized into animal tissues, while carbon can be metabolically routed to either direct incorporation to tissue or to storage within large lipid stores depending on feeding status (69). Under fasting conditions, lipid stores will be utilized as a source of C with a light δ^13^C value that reflects the fractionation of C from food resources containing the regional δ^15^N values where they were consumed (70). These lipid stores are expected to be mobilized during times of limited feeding and reduce the impact of C derived from incidental feeding on δ^13^C_Base_ values of the baleen during fasting conditions as metabolites from blood are incorporated into baleen. This metabolic routing of C would contribute to masking the variation in C values along the baleen from incidental feeding. δ^15^N_Base_ values (range of minimums from 2.8 to 5.8‰) never reach the expected low isoscape values for the mid-Atlantic of ∼ −1 to 1‰ as the lower concentrations of N from opportunistic feeding may be insufficient to fully turnover the comparatively high δ^15^N fed on extensively at higher latitude feeding grounds (6 to 10‰.).

Differentiation between periods of migration is supported in four individuals (FW1-FW3, MN), with clear separation between niche areas for basal resources that predominantly separate along the δ^15^N_Base_ axis (Overlap < 27%, Hotelling’s T^2^-test, *p*<0.001; Fig. 5). The humpback whale appears to predominately separate along the δ^13^C_Base_ axis which may reflect utilization of benthic resources within coastal margins, a lack of opportunistic feeding during migrations (17), or a relatively reduced latitudinal migration indicative of a temperate feeding style where individuals preferably feed in the temperate zone and reduce reliance on high-latitude feeding (53). The gradual increase in δ^15^N value across the baleen record of the humpback whale from around 11.5 to 15‰ likely reflects increased contribution of herring and other small fish during feeding. High δ^15^N_Gly_ values observed for the humpback (∼0‰; Fig. 4b) also mirror what is expected for coastal predation on small fish migrating from estuarine into coastal waters where elevated δ^15^N_Gly_ values have been previously observed such as herring (42).

In the above discussions it should be noted that individuals FW1 and FW3 have statistically relevant differences between δ^15^N_Phe-Base_ and δ^15^N_Gly_, were males, and the remaining three individuals were female. Females are likely to display different isotopic patterns for both C and N as they reproduce, as the result of gestation and lactation altering the partitioning of resources and resultant isotopic values (71). These effects remain unaccounted for in this analysis due to the limited knowledge of these individual’s life histories as baleen was sampled from beaching and bow-catch events. Additionally, the use of isoscapes to provide yearly averages for underlying isotope values for POM ignores variability that is expected during seasonal changes, although examinations of this variance are increasingly common (54, 72). Seasonal and temporal isotopic variations can be quite large (35, 68), especially for arctic or near arctic waters where the bulk of mysticete feeding occurs, and should be further considered in future work.

## Conclusion

This study utilizes isotope analysis of bulk material and amino acids to provide metabolic and trophic context to the isotopic oscillations observed within whale baleen. Trophic level estimates from source and trophic amino acids were higher for the humpback and minke whales than for the fin whales, which corresponds to previous observations. Trophic level estimates using amino acids allow for the correction of bulk stable isotope values from consumers to the underlying baseline values of the primary producers supporting regional foodwebs. Isotopic niche areas constructed from these baseline values using periods of minimum and maximum bulk δ^15^N values further confirm distinct differences between maximum periods that reflect feeding in high-latitude feeding grounds and minimum periods that reflect resources utilized from mid-Atlantic breeding grounds for these individuals. δ^15^N values from Gly and Thr provided useful metabolic indicators across the baleen sequences. Gly had higher values that aligned with considerable time spent in breeding grounds where feeding is expected to become incidental and fasting is likely to occur. Differences in Thr occurred more episodically and indicate that food stress or starvation occurred more often in the fin whales in this study than in the humpback or minke whales. Analysis of δ^15^N values for amino acids provided further context into the movements and metabolic condition for mysticete whales, information that is especially important in wide ranging, difficult to track, animals with threatened conservation status.

## Acknowledgements

We thank Jort Ossebaar, Monique Verweij, and Ronald von Bommel for technical support for this project. Funding was provided through the project “Waddensleutels” funded by “Waddenfonds” (WF203930). We thank Kees Camphuysen for the input provided during the conceptual stages of this work. Clive Trueman provided considerable food for thought on the first draft of this manuscript and we are grateful for his considered input and conversation during lockdown.

## Supplementary Material

**Supplemental table 1:**

| Species | ID | Samples<br>(n) | $\delta^{13}\text{C}$ (‰) | | | $\delta^{15}\text{N}$ (‰) | | |
| --- | --- | --- | --- | --- | --- | --- | --- | --- |
|  |  |  | Max | Mean | Median | Max | Mean | Median |
| | | | Min | (±SD) | | Min | (±SD) | |
| Fin whale | 1 | 43 | -18.4<br>-19.3 | -18.8<br>(0.2) | -18.9 | 10.6<br>9 | 9.8<br>-0.4 | 9.9 |
|  | 2 | 36 | -18.9<br>-19.5 | -19.2<br>(0.9) | -19.1 | 10.7<br>8.9 | 9.9<br>-0.5 | 10 |
|  | 3 | 105 | -17.9<br>-20 | -18.9<br>(0.5) | -18.9 | 9.9<br>7.1 | 9.1<br>-0.6 | 9.3 |
| Humpback whale |  | 27 | -19.1<br>-19.9 | -19.6<br>(0.2) | -19.6 | 14.8<br>11.1 | 12.8<br>-1.1 | 12.8 |
| Minke whale |  | 48 | -17.6<br>-18.9 | -18.1<br>(0.3) | -18.1 | 12.9<br>10 | 11.8<br>-0.8 | 12.2 |

**Supplemental Table 2:**
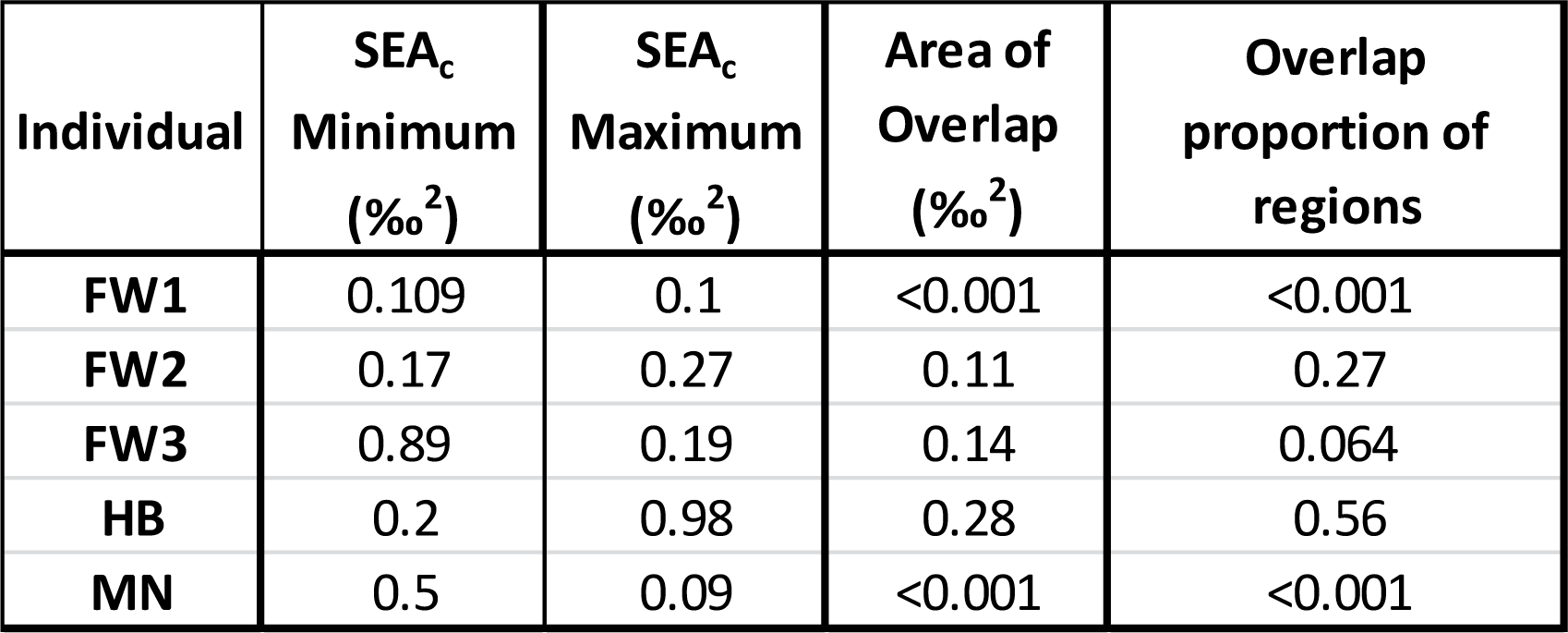
Standard ellipse areas and overlap proportions for minimum and maximum baleen regions.

